# Gut microbiome and its cofactors are linked to lipoprotein distribution profiles

**DOI:** 10.1101/2021.09.01.458531

**Authors:** Josué L. Castro-Mejía, Bekzod Khakimov, Mads V. Lind, Eva Garne, Petronela Paulová, Elnaz Tavakkoli, Lars H. Hansen, Age K. Smilde, Lars Holm, Søren B. Engelsen, Dennis S. Nielsen

## Abstract

Increasing evidence indicates that the gut microbiome (GM) plays an important role in the etiology of dyslipidemia. To date, however, no in-depth characterization of the associations between GM and its metabolic attributes with deep profiling of lipoproteins distributions (LPD) among healthy individuals has been conducted. To determine associations and contributions of GM composition and its cofactors with distribution profiles of lipoprotein subfractions, we studied blood plasma LPD, fecal short-chain fatty acids (SCFA) and GM of 262 healthy Danish subjects aged 19-89 years.

Stratification of LPD segregated subjects into three clusters of profiles that reflected differences in the lipoprotein subclasses, corresponded well with limits of recommended levels of main lipoprotein fractions and were largely explained by host characteristics such as age and body mass index. Higher levels of HDL, particularly driven by large subfractions (HDL2a and HDL2b), were associated with a higher relative abundance of Ruminococcaceae and Christensenellaceae. Increasing levels of total cholesterol and LDL, which were primarily associated with large 1 and 2 subclasses, were positively associated with Lachnospiraceae and Coriobacteriaceae, and negatively with Bacteroidaceae and Bifidobacteriaceae. Metagenome sequencing showed a higher abundance of genes involved in the biosynthesis of multiple B-vitamins and SCFA metabolism among subjects with healthier LPD profiles. Metagenomic assembled genomes (MAGs) affiliated mainly to Eggerthellaceae and Clostridiales were identified as the contributors of these genes and whose relative abundance correlated positively with larger subfractions of HDL.

The results of this study demonstrate that remarkable differences in composition and metabolic traits of the GM are associated with variations in LPD among healthy subjects. Findings from this study provide evidence for GM considerations in future research aiming to shade light on mechanisms of the GM – dyslipidemia axis.

## INTRODUCTION

Cholesterol is essential for keeping cellular integrity and is an important precursor for steroid hormones and bile acids ^1^. However, alterations of the cholesterol metabolism and consequent dyslipidemia have been associated with various diseases, including atherosclerosis and cardiovascular diseases (CVD) ^2^, as well as breast cancer ^3^.

Recent advances in metabolomics research have allowed large-scale and high-throughput profiling of lipoprotein distribution’s (LPD) in human blood plasma based upon their composition and concentration ^4–6^. It has been hypothesized that numerous medical conditions such as glucose intolerance, type-2 diabetes, myocardial infarction, ischemic stroke and intracerebral hemorrhage, might be associated with lower blood levels of larger HDL particles (e.g. HDL2a and HDL2b) and a higher content of triglycerides within the lipoproteins particles ^7,8^.

During the last decade it has been shown that alterations in gut microbiome (GM) composition contribute to the development and progression of several metabolic and immunological complications ^9^. Furthermore, a handful of recent studies on different cohorts have also demonstrated that the changes in intestinal microbiota are highly correlated to variations in levels of lipoproteins in blood ^10–12^, as well as to promote atherosclerosis ^13^, and regulate cholesterol homeostasis ^14^.

The relationship between GM and LPD has only been scarcely investigated. Recently Vojinovic et al. ^5^ reported the association of up to 32 GM members with very-low-density (VLDL) and high-density (HDL) subfractions. Positive correlations between a number of Clostridiales members with large particle size subfractions of HDL were elucidated. In other studies, focusing on total lipoproteins fractions, an increasing abundance of GM members affiliated to the Erysipelotrichaceae and Lachnospiraceae families have been linked to elevated levels of total cholesterol and low-density lipoproteins (LDL) ^10–12^. Interestingly, common gut microbes like Lactobacillaceae members have been reported to assimilate and lower cholesterol concentrations from growth media and incorporate it into their cellular membrane ^15^, whereas butyrate-producing *Roseburia intestinalis* has been found to increase fatty acid utilization and reduce atherosclerosis development in a murine model ^16^.

However, the relationship between GM and LPD distribution is still far from being understood. Thus, with the aim of gaining a deeper understanding of the relationship between GM and LPD in blood, we carried out a detailed compositional analysis of GM, its metabolic functions, and studied its associations with blood lipoproteins quantified using a recently developed method based on proton (^1^H) nuclear magnetic resonance (NMR) spectroscopy ^6^. We determined covariations between larger HDL subclasses and lower total cholesterol with a several Clostridiales (Ruminococcaceae and Lachnospiraceae) and Eggerthelalles members, whose metabolic potential is linked to biosynthesis of cofactors essential for carrying out lipid metabolism.

## METHODS

### Study participants

Two hundred and sixty-two men and women participants older than 20 years, who had not received antibiotic treatment 3 months prior to the beginning of the study and who had not received pre- or probiotics 1 month prior to the beginning of the study, were included as part of the COUNTERSTRIKE (COUNTERacting Sarcopenia with proTeins and exeRcise – Screening the CALM cohort for lIpoprotein biomarKErs) project (counterstrike.ku.dk). Pregnant and lactating women, as well as individuals suffering from CVD, diabetes or chronic gastrointestinal disorders, were excluded from the study.

### Ethics approval and consent to participate

The study was approved by the Research Ethics Committees of the Capital Region of Denmark in accordance with the Helsinki Declaration (H-15008313) and the Danish Data Protection Agency (2013-54-0522). Written informed consent was obtained from all participants.

### Lipoprotein distribution profiles

The human blood plasma lipoproteins were quantified using SigMa LP software ^17^. The SigMa LP quantifies lipoproteins from blood plasma or serum using optimized partial least squares (PLS) regression models developed for each lipoprotein variable using one-dimensional (1D) ^1^H NMR spectra of blood plasma or serum and ultracentrifugation based quantified lipoproteins as response variables as determined in Khakimov et al. ^6^.

### Short chain fatty acids (SCFAs) quantification

Targeted analysis and quantification of SCFA on fecal slurries were carried out as recently described ^18^

### Samples processing, library preparation and DNA sequencing

Fecal samples were collected and kept at 4°C for maximum 48 h after voidance and stored at −60°C until further use. Extraction of genomic DNA and library preparation for high-throughput sequencing of the V3-region of the 16S rRNA gene was performed as previously described ^18^. Shotgun metagenome libraries for sequencing of genome DNA were built using the Nextera XT DNA Library Preparation Kit (Cat. No. FC-131-1096) and sequenced with Illumina HiSeq 4000 by NXT-DX.

### Analysis of sequencing data

The raw dataset containing pair-end amplicon reads we analyzed following recently described procedures ^18^. The metabolic potential of the amplicon sequencing dataset was determined through PICRUSt ^19^, briefly, zero-radious operational taxonomical units (zOTUs) abundances were first normalized by copy number and then KEGG orthologues was obtained by predicted metagenome function.

For shotgun sequencing, the reads were trimmed from adaptors and barcodes and the high-quality sequences (>99% quality score) using Trimmomatic v0.35 ^20^ with a minimum size of 50nt were retained. Subsequently, sequences were dereplicated and check for the presence of Phix179 using USEARCH v10 ^21^, as well as human and plant genomes associated DNA using Kraken2 ^22^. High-quality reads were then subjected to within-sample *de-novo* assembly-only using Spades v3.13.1 ^23^ and the contigs with a minimum length of 2,000 nt were retained. Within-sample binning was performed with metaWRAP ^24^ using Metabat1 ^25^, Metabat2 ^26^ and MaxBin2 ^27^, and bin-refinement ^28^ was allowed to a ≤10% contamination and ≥70% completeness. Average nucleotide identity (ANI) of metagenome bins, or metagenome assembled genomes (MAGs), was calculated with fastANI ^29^ and distances between MAGs were summarized with bactaxR ^30^. To determined abundance across samples, reads were mapped against MAGs with Subread aligner ^31^ and a contingency-table of reads per Kbp of contig sequence per million reads sample (RPKM) was generated. Taxonomic annotation of MAGs was determined as follows: ORF calling and gene predictions were performed with Prodigal ^32^, the predicted proteins were blasted (blastp) against NCBI NR bacterial and archaeal protein database. Using Basic Sequence Taxonomy Annotation tool (BASTA) ^33^, the Lowest Common Ancestor (LCA) for every MAG was estimated based on percentage of hits of LCA of 60, minimum identity of 0.7, minimum alignment of 0.7 and a minimum number of hits for LCA of 10.

To determine the metabolic potential of metagenomes, ORF calling and gene predictions (similar as above) were performed on both, binned and unbinned contigs, and the predicted proteins were subsequently clustered at 90% similarity using USEARCH v10. To assign functions, protein sequences were blasted (90% id and 90% cover query) against the integrated reference catalog of the human gut microbiome (IRCHGM) ^34^, while using only target sequences containing KEGG ortholog entries. Similar as above, to determine abundance of protein-encoding genes across metagenomes, reads were mapped against protein clusters (PC) with DIAMOND ^35^ and a contingency-table of reads mapped to PCs was also generated. To avoid bias due to sequencing depth across protein-encoding genes, samples were subsampled to 15,000,000 reads per sample.

### Statistical analysis

Stratification and clustering of LPD was carried out using Euclidean distances and general agglomerative hierarchical clustering procedure based on “Ward2”, as implemented in the *gplots* R-package ^36^. For univariate data analyses, pairwise comparisons were carried out with unpaired two-tailed Student’s *t*-test, Spearman’s rank coefficient was used for determining correlations and Chi-Square test for evaluating group distributions. For multivariate data analyses, the association of covariates (e.g. age, BMI, sex) with LPD were assessed by redundancy analysis (RDA) (999 permutations), whereas the association of LPD clusters with GM were analyzed by distance-based RDA (999 permutations) on Canberra distances (implemented in the *vegan* R-package ^37^).

Feature selection for zOTUs was performed with Random Forest. Briefly, for a given training set (training: 70%, test: 30%), the *party* R-package ^38^ was run for feature selection using unbiased-trees (cforest_unbiased with 6,000 trees and variable importance with 999 permutations) and subsequently the selected variables were used to predict (6,000 trees with 999 permutations) their corresponding test set using *randomForest* R-package ^39^. The selected features were subjected to sequential rounds of feature selection until prediction could no longer be improved. All statistical analyses were performed in R versions ≤3.6.0.

### Data availability

Sequence data are available at the Sequence Read Archive (SRA), BioProject SUB9304449 submissions SUB9305011 and SUB9304442. Supplementary Table 1 provides samples information. Non-sequence data that support the findings of this study are available from the corresponding authors upon reasonable request.

## RESULTS

### Participants and data collection

Two hundred and sixty two individuals (men:women 90:172) with an age between 20 and 85 years (Figure 1A) and BMI ranging between 19 and 37 kg/m^2^ (Figure 1B) were included in this study. Subjects are representatives of community dwelling and apparently healthy adults living in the Danish Capital Region. In this study, we included ^1^H NMR spectroscopy based quantified lipoproteins from human blood plasma^6^, short-chain fatty acids profiling and GM composition on fecal samples based on 16S rRNA-gene amplicon sequencing and shotgun metagenome sequencing for a subset of samples (Figure 1C).

**Figure 1.**
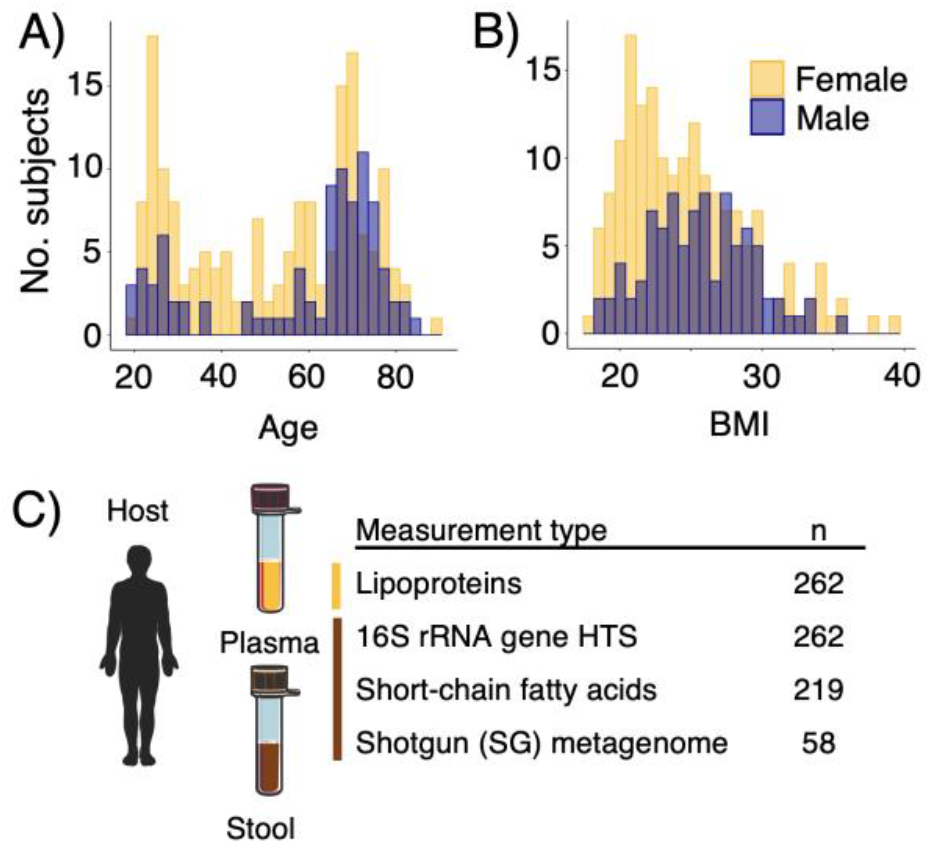
COUNTERSTRIKE participants and sample overview. **A**) Age and **B**) body mass index (BMI) distribution of the study participants. **C**) samples and datasets included and analyzed in this study.

### LPD profiles, stratification and host covariates

LPD profiles of the study subjects were predicted from ^1^H NMR measurements of blood plasma. A total of 55 lipoproteins-subfractions were quantified including cholesterol, triglycerides (TG), cholesterol ester (CE), free cholesterol, phospholipids, apolipoprotein A (ApoA1) and apolipoprotein B (ApoB) content in all or in some of lipoprotein in plasma (VLDL, IDL, HDL, LDL) and/or in lipoprotein subfractions (HDL2a, HDL2b, HDL3, LDL1, LDL2, LDL3, LDL4, LDL5, LDL6)^6^. Linking host covariates and LPD profiles, redundancy analysis (RDA) of LPD profiles showed a significant (*p* ≤ 0.01) effect of age, BMI and sex on LPD profiles (Figure 2B) with a combined size effect of up to 24.6% (Figure 2B-C).

**Figure 2.**
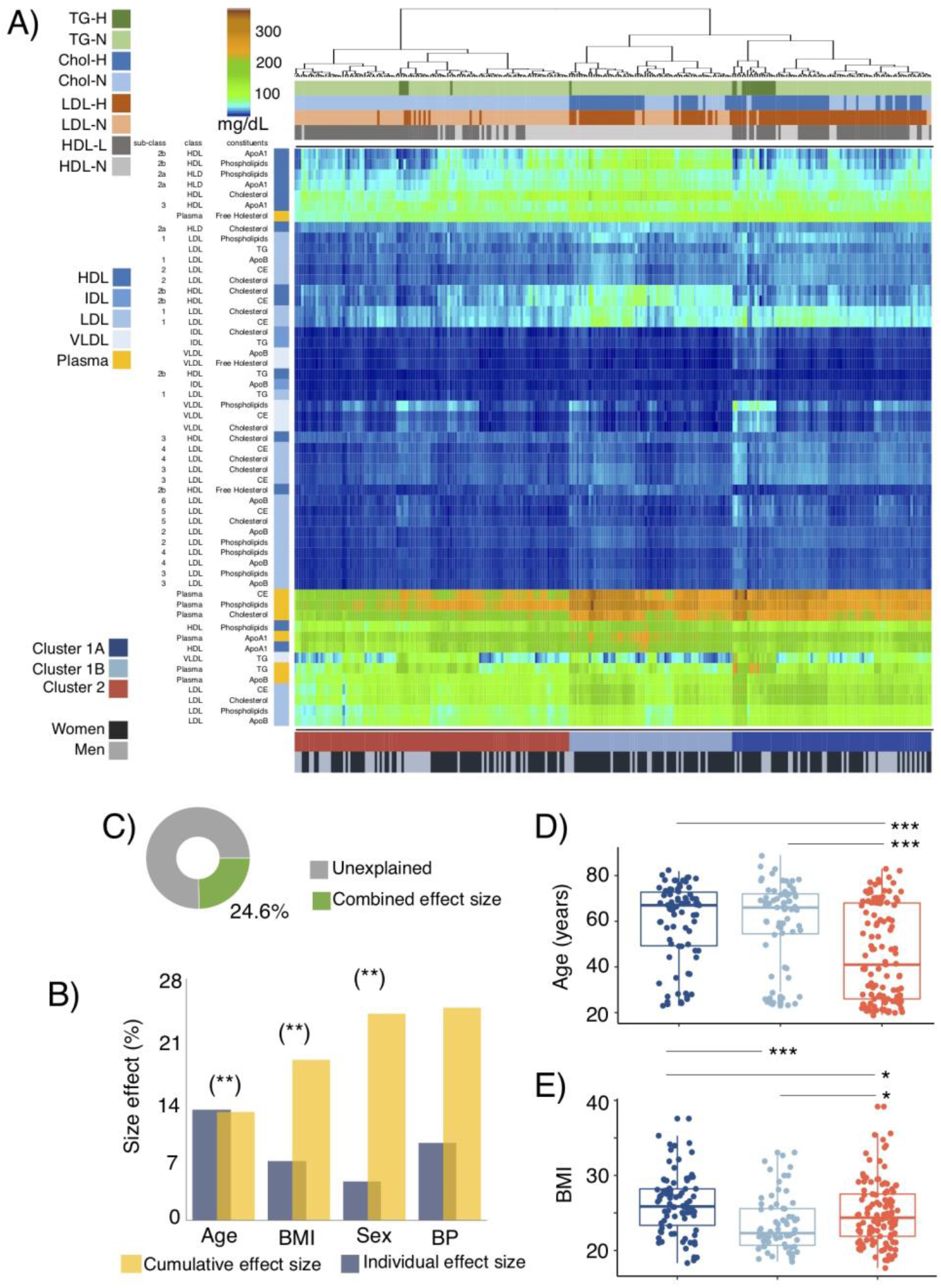
Plasma lipoprotein distribution (LPD) profiles and covariates. **A**) Profiles of main and sub-fractions of plasma lipoprotein distribution (LPD) determined by ^1^H-NMR^6^. LPD are clustered using Euclidean distances and general agglomerative hierarchical clustering procedure. Upper color bars represent within-/out-of the recommended levels of main lipoprotein fractions suggested by the NIH ^40^ (total cholesterol <200mg/dL, LDL <100mg/dL, HDL >60mg/dL, Triglycerides <150 mg/dL). Lower color bars depict 3 clusters (C1A, C1B and C2) of study participants given their LPD profile and the sex distribution of subjects. **B**) Cumulative effect size of non-redundant covariates of LPD determined by stepwise RDA analysis (right bars) as compared to individual effect sizes assuming independence (left bars). **C**) Fraction of LPD variation explained with the stepwise approach. Distribution of **D**) age and **E**) body mass index (BMI) between subjects belonging to C1A, C1B and C2. Stars show statistical level of significance (\**p*≤ 0.05, \*\**p*≤ 0.01, \*\*\**P*≤ 0.001)

Clustering of LPD profiles segregated study participants into three groups (Figure 2A, Figure I in the Data Supplement). Cluster 1A and 1B were characterized by higher concentrations of LDL sub-fractions and their constituents (particularly evident in subclasses 1 and 2). Clusters 1A and 2, on the other hand, were characterized by lower concentrations of HDL sub-fractions (associated with HDL2a and HDL2b), whereas higher concentrations of HDL-3 particles in subjects of cluster 1A were observed (Figure I in the Data Supplement). Furthermore, plasma concentrations of CE, phospholipids and CE were higher among cluster 1A and 1B. When comparing the plasma fractions of the study participants to the recommendations of cholesterol classes provided by the National Institute of Health (NIH) ^40^, for clusters 1A and 1B total cholesterol and LDL levels were above the recommendations, while for clusters 1B and 2 the levels pf HDL were below the recommended values.

LPD profiles were also found to covariate with host attributes, cluster 2 subjects was significantly younger than clusters 1A and 1B (Figure 2D), and cluster 1B showed the lowest BMI (Figure 2E). These results were also consistent even after correcting for sex effects, given that cluster 1B had a significantly higher proportion of women (Fisher test *p* < 0.01, Figure 2A) compared to clusters 1A and 2 (Figure I in the Data Supplement).

### LPD clusters are linked with GM profiles

The GM of study participants (n = 262) was profiled using high-throughput amplicon sequencing the V3-region of the 16S rRNA gene (11,544 zOTUs), as well as shotgun metagenome sequencing of total genomic DNA for a subset of samples (n = 58). Gene content and functionality (based on KEGG orthologues - KOs) were predicted based on PICRUSt ^19^ (for 16S rRNA gene amplicons), as well as through ORF calling and gene prediction of assembled contigs reconstructed from shotgun metagenome data. Validation of PICRUSt against metagenome calling KO yielded a high correlation coefficient (Pearson *r* = 0.77, Figure 3A) between the gene richness of both datasets. Alpha diversity analyses between LPD clusters revealed no significant (*t-* test *p* > 0.05) differences in phylotypes (Figure 3B) nor KOs richness as predicted by the PICRUSt (Figure 3C). A significant (Dip-test *p* < 0.001) bimodal distribution of KO richness among the study participants was observed, but a higher-/lower-gene count was not associated to LPD clusters (Figure 3C) or BMI categories (Figure 3D). Significant differences in composition (beta-diversity) between LPD clusters were observed among phylotypes (Canberra distance, Adonis test *p* < 0.05, R^2^ = 0.62-1%), but not among PICRUSTs predicted KOs.

**Figure 3.**
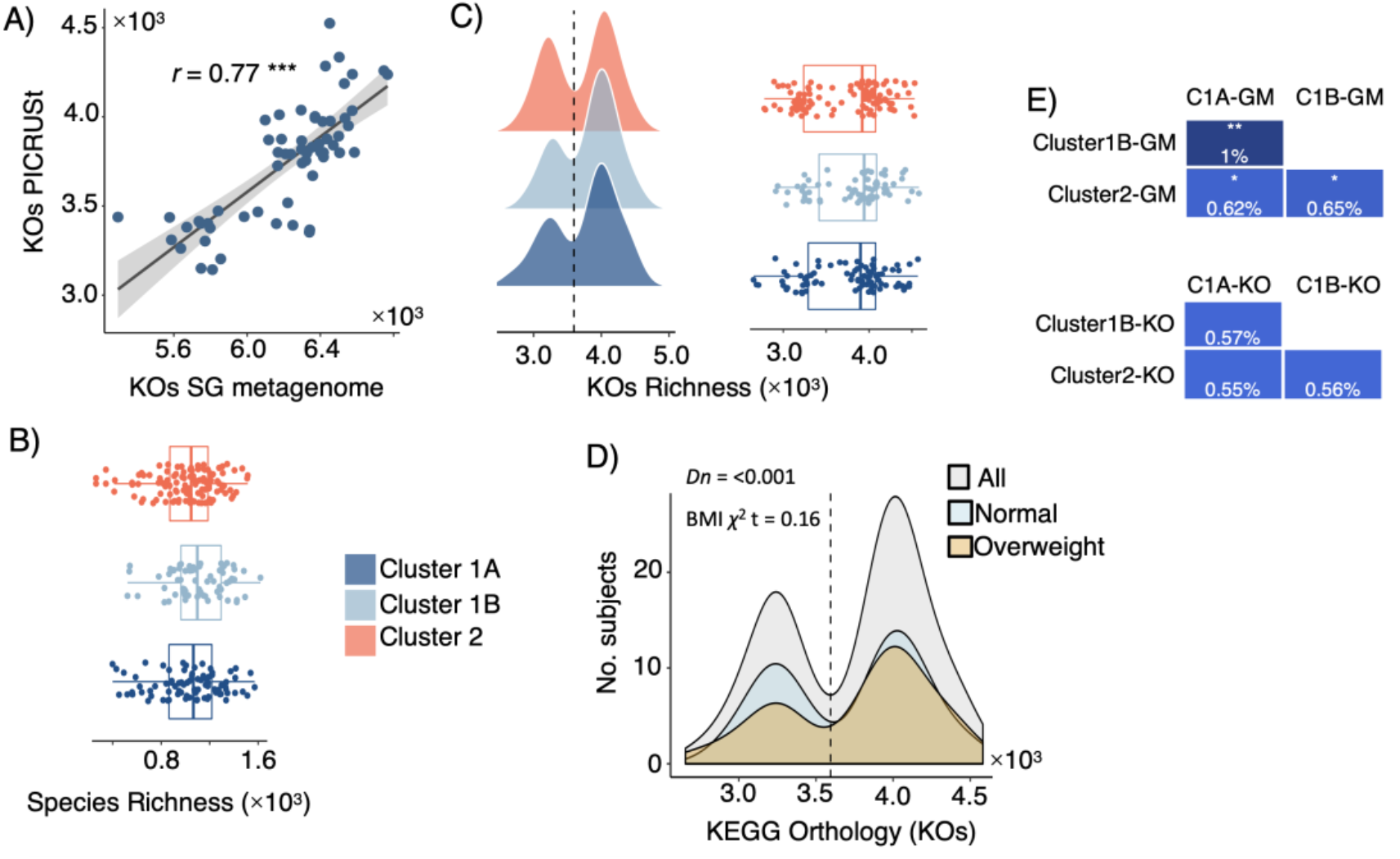
Diversity metrics on gut microbiota and metabolic content. **A**) Spearman’s rank correlation between fecal microbial KEGG Orthologues (KOs) from shotgun metagenome (SG) sequencing and KO predicted by PICRUSt. **B**) Richness of microbial phylotypes (zOTUs) richness and **C**) KO predicted by PICRUSt among subjects catalogued as being C1A, C1B and C2 based on their LPD. **D**) KO counts (richness) among all subjects and those with BMI ≤ 25 (normal) and BMI >25 (overweighed); the observed bimodal distribution was statistically significant by the dip-test. **E**) Adonis test based on Canberra dissimilarities quantifying variance explained (*R*^2^) and significance of phylotypes and KO abundance with LPD clustering. Stars show statistical level of significance (\**p*≤ 0.05, \*\**p*≤ 0.01, \*\*\**P*≤ 0.001)

### LPD clusters correspond with GM and KOs features

After feature selection based on random forest, LPD clusters were partially discriminated (Figure 4A) by 206 selected sequence variants (zOTUs) distributed to over 10 families (Figure 4B). Among these, zOTUs affiliated to Ruminococcaceae (75) and Lachnospiraceae (58) represented 64%, followed by Bacteroidaceae (8), Bifidobacteriaceae (7), Christensenellaceae (6), Coriobacteriaceae (5) and four other sparse bacterial families (47). The cumulative abundance (cumulative sum scaling, CSS) of those families showed differences between LPD clusters, with cluster 1A being associated with a higher abundance of Lachnospiraceae and a lower abundance of Christensenellaceae members, while cluster 1B was characterized by a larger proportion of Ruminococcaceae phylotypes, and cluster 2 showed increased proportion of Bifidobacteriaceae, Bacteroidaceae and reduced abundance of Coriobacteriaceae (Figure 4B-C).

**Figure 4.**
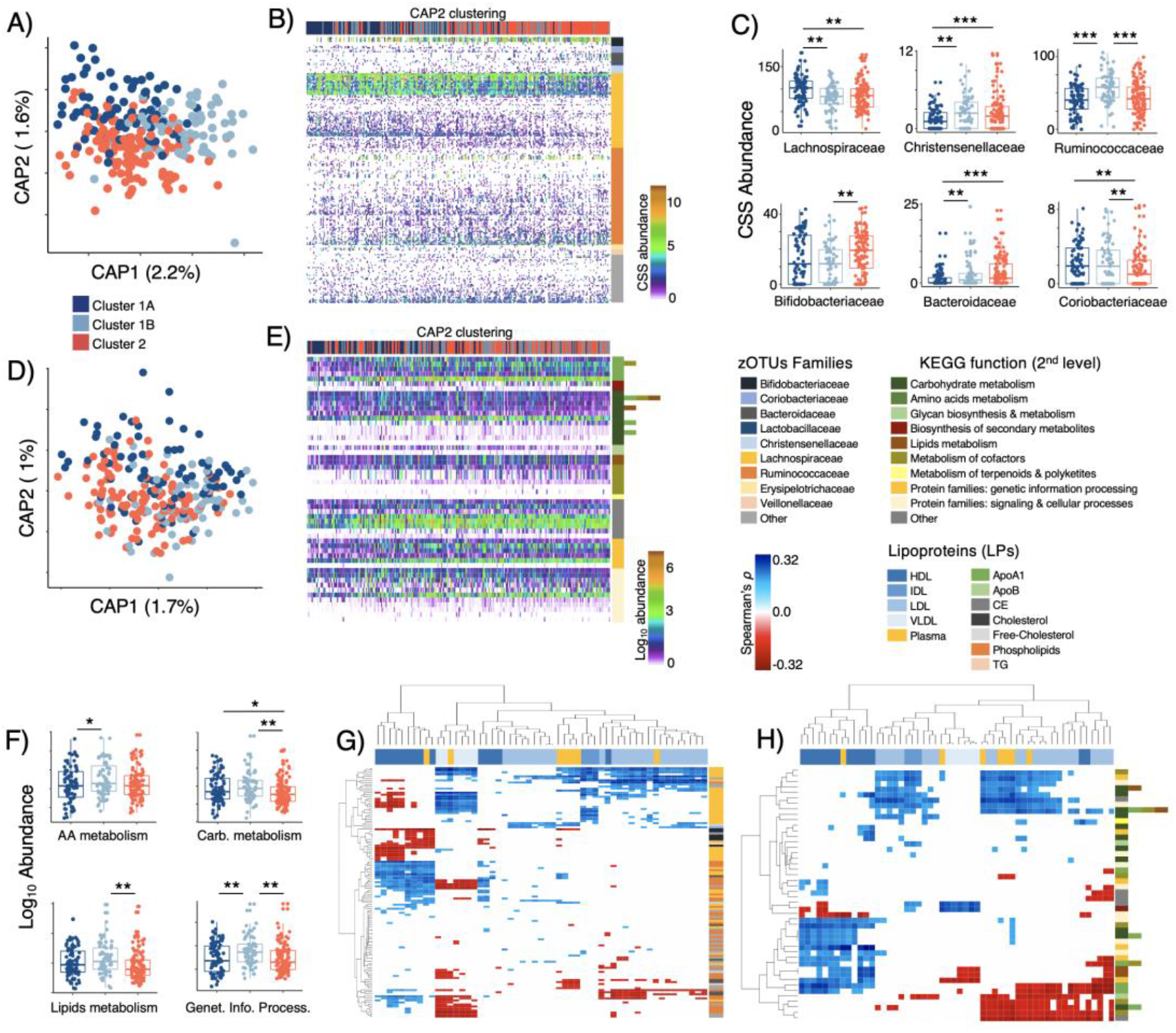
Phylotypes and KO functions associated with LPD clustering. Distance-based RDA (Canberra dissimilarity) displaying discrimination of LPD clusters based on selected **A**) zOTUs (*p* = 0.001, explained variance = 3.8%) and **D**) KOs-PICRUSt (*p* = 0.001, explained variance = 2.7%) selected through Random Forests. Overview of selected **B**) zOTUs and **E**) KOs-PICRUSt clustered using Canberra distances and general agglomerative hierarchical clustering procedure based on ward2. Distribution of **C**) zOTUs summarized to family level and **F**) KOs-PICRUSt summarized to 2^nd^ level KEGG function across subjects belonging C1A, C1B and C2 LPD groups. Heatmaps displaying significant (False Discovery Rate corrected, FDR ≤ 0.05) Spearman’s rank correlations between **G**) zOTUs and LPD sub-fractions, as well as **H**) KOs-PICRUSt and LPD sub-fractions. Stars show statistical level of significance (\**p*≤ 0.05, \*\**p*≤ 0.01, \*\*\**P*≤ 0.001)

KEGG orthologues predicted through PICRUSt demonstrated very weak discrimination power towards LPD clusters (Figure 4D, Figure II-A in the Data Supplement shows detailed 3^rd^ level KEGG functions), this included 54 KOs affiliated to >9 primary and secondary metabolism processes, as well as signaling and cellular processes (Figure 4E). Despite its documented limitations ^41^ PICRUSt was still able to reveal a decreasing abundance of functional modules among subjects of cluster 1A and 2 as compared to those of cluster 1B (Figure 4E-F). Analysis on aggregated functions per KOs (2^nd^ level KEGG) showed that cluster 1B was characterized by a higher abundance (t-test *p* < 0.05) of functions related to metabolism of amino acids (e.g., Phe, Tyr and Trp biosynthesis), carbohydrates (e.g., pyruvate, propanoate and butanoate metabolism), lipids (glycerolipids and glycerophospholipids metabolism) and genetic information processing (e.g., transcriptional factors) (Figure 4F).

Correlation analyses of selected zOTUs vs LPD profiles displayed several significant (Spearman FDR p ≤ 0.05) associations (Figure 4G, Figure II-B in the Data Supplement). Most Ruminococcaceae (74/75 phylotypes, mostly unclassified), a division of Lachnospiraceae (13/58 phylotypes, mostly unclassified), Bacteroidaceae (e.g., *B. massiliensis*, *B. caccae*) Christensenellaceae (unclassified genus) and Coriobacteriaceae (unclassified genus) showed positive correlations with HDL subfractions and negative correlations with VLDL and LDL (e.g. LDL3, 4, 5, 6). Contrary to this, most Lachnospiraceae (45/58), Veillonellaceae (e.g., *V. invisus*) and Bifidobacteriaceae (e.g., *Bf. adolescentis*, *Bf. bifidum*) phylotypes correlated negatively with HDL subfractions, and positively with subfractions composed of IDL, LDL and VLDL. For KOs vs LPD (Figure 4H, Figure II-C in the Data Supplement), increasing abundance of functions linked to glycerophospholipids metabolism and amino acids (His, Phe, Tyr and Trp) biosynthesis correlated positively with HDL fractions and negatively with LDL and VLDL. Furthermore, the production of glycosphingolipids, biotin (Vit_B7_) and lipopolysaccharides correlated negatively with small LDL subfractions (e.g. LDL3, 4, 5, 6).

### Metagenome bins and functions associated with LPD clusters

Fifty-eight samples were subjected to shotgun metagenome sequencing (Figure 1C) generating on average 5.2 GB per sample. ORF calling on the entire assembled dataset of generated ~1.4 million gene-clusters (90% similarity clusters, here termed “genes”), with 84,560 core genes being present in at least 90% of the metagenome sequenced samples. RDA analysis of the core-gene dataset showed significant (*p* = 0.001) differences between LPD clusters and explaining up to 23.7% of the total variance in gene composition (Figure 5A). Ranking of variables (i.e. top 150) within the 1^st^ and 2^nd^ canonical components of the CAP analyses provided an overview of 35 “known” metabolic genes (>90% identity match to the integrated non-redundant gene catalog with KEGG ortholog entries ^34^, Figure 5B, Figure III-A in the Data Supplement) linked to >10 2^nd^ level KEGG functions, which resembled the large majority of those predicted by PICRUSt (see Figure 4E-F). A higher abundance of these genes was observed among subjects grouped within Cluster 1B relative to cluster 1A and Cluster 2. To determine the species associated with these genes, gene-sequences were mapped back to 1,419 metagenome-assembled genomes (MAGs) (Figure 5C). Sixty MAGs affiliated to Lachnospiraceae, Clostridiales, Coriobacteriaceae and Firmicutes and clustered within 19 species were found to contribute with 27 out of the 35 genes that discriminated LPD clusters (Figure 5D, Figure III-B in the Data Supplement). MAGs-G1 to G5 contributed with peptidoglycan and glycan biosynthesis. MAGs-G6 to G12 contributed with thiamine (Vit_B1_) and pantothenate (Vit_B5_) metabolism, starch degradation and butyric acid metabolism (butanol dehydrogenase that may lead to increased concentrations of 1-butanol at the expense of butyrate production, Figure 5E) and glycerolipid metabolism. Finally, MAGs-G13 to G19 promoted biosynthesis of glucosinates, metabolism of propionic acid, biosynthesis of fatty acids, Vit_B6_ metabolism, as well as folate (Vit_B9_) biosynthesis (Figure 5D, 5F, Figure III-B in the Data Supplement). Subjects belonging to LPD-cluster 1B had a significantly higher relative abundance of MAGs-G7, MAGs-G9 to G19 (those comprising Clostridiales, Eggerthellaceae and Firmicutes bins, Figure 5G-H), MAGs-G1 and MAGs-G5 (those affiliated to Lachnospiraceae, Figure 5I) than subjects in clusters 1A and 2. Likewise, their cumulative abundance reached significant positive (spearman *p* < 0.001) correlations with constituents (e.g., Cholesteryl ester) of larger HDL sub-classes (HDL2a and HDL2b) (Figure 5J).

**Figure 5.**
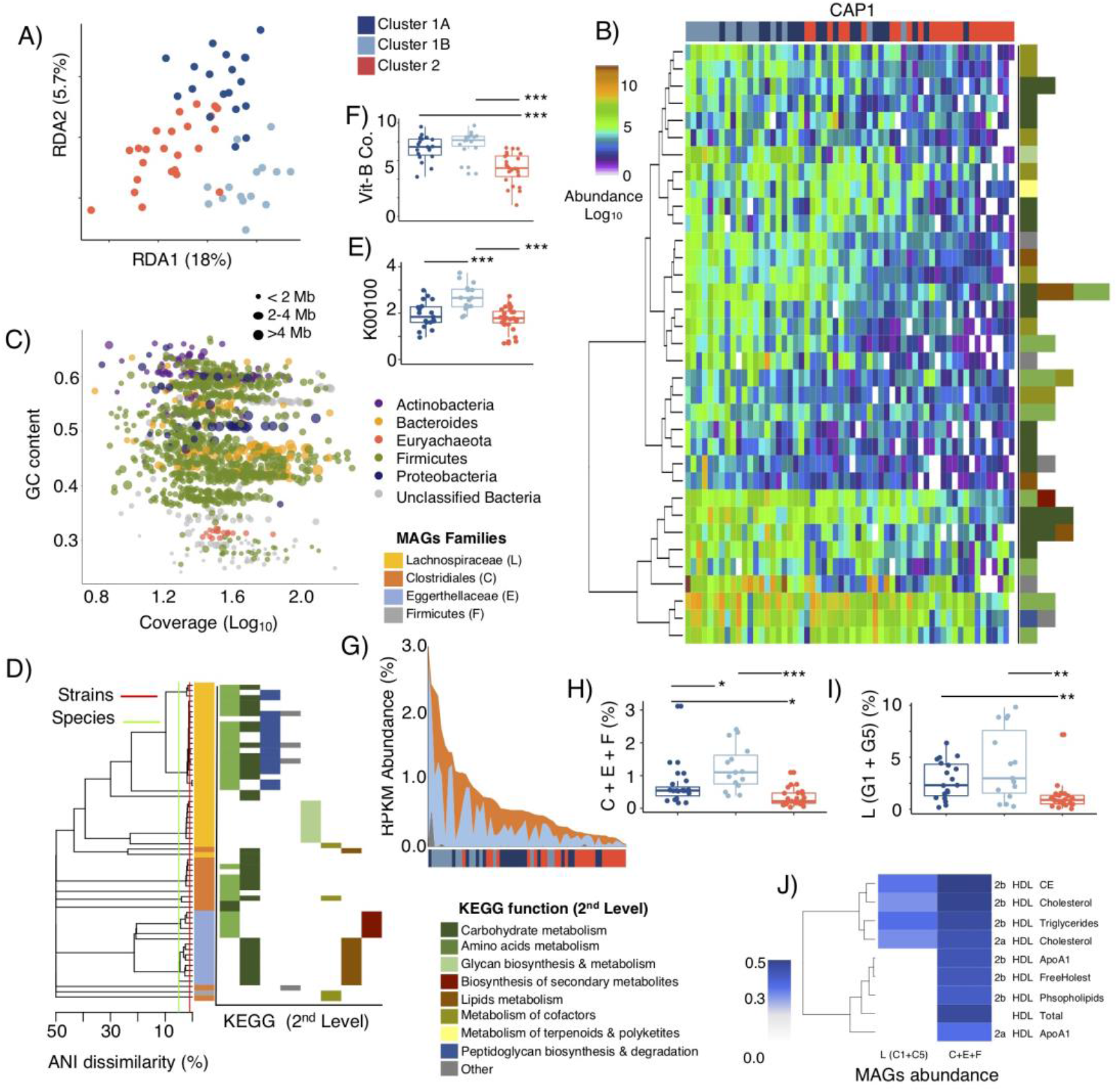
Metagenome metabolic functions and associated MAGs. **A**) RDA displaying discrimination of LPD clusters based on selected KOs obtained from shotgun metagenome and assembly (*p* = 0.001, explained variance = 23.7%). **B**) Overview of most discriminatory (based on CAP1 and CAP2 within db-RDA) KOs with known metabolic functions clustered using Canberra distances and general agglomerative hierarchical clustering procedure based on ward2. **C**) GC-content – Coverage plot of metagenome assembled genomes (MAGs) with ≤10% contamination and ≥70% completeness. MAGs are colored according to phylum-level taxonomic affiliation and bubble size indicates their genome size in mega-bases (Mb). **D**) Phylogeny of MAGs containing KOs that discriminate LPD clusters (1A, 1B and 2), a cut-off of 95-ANI (species-level) and 99-ANI (strain-level) are denoted. MAGs are colored at family level affiliations and their KOs contribution at the 2^nd^ level KEGG function pathways are provided. **E**) Relative abundance of protein-encoding genes associated with butanol dehydrogenase (K00100), and **F**) protein-encoding genes associated metabolism and biosynthesis of vitamin B1, B2, B5 and B9. **G-H**) Distribution of cumulative abundance (RPKM) of MAGs (containing discriminatory KOs) associated with Clostridiales, Coriobacteriaceae and Firmicutes (Cl + Co + F) among LPD clusters. **I**) Distribution of cumulative abundance (RPKM) of MAGs (G1 + G5 – see Figure III-B in the Data Supplement, containing discriminatory KOs) associated with Lachnospiraceae among LPD clusters. **I**) Heatmaps displaying significant (False Discovery Rate corrected, FDR ≤ 0.05) Spearman’s rank correlations between MAGs and HDL subfractions.

The concentrations of the SCFAs acetate and propionate in fecal samples showed no differences between LPD clusters. However, higher concentrations of butyrate, isobutyrate, 2-methylbutyrate, valerate and isovalerate (ANOVA Tukey’s HSD *p* < 0.05) were observed in cluster 2 (Figure 6A-D). To determine whether microbial activity was linked to the production of such branched-chain fatty-acids, we then focused on analyzing the abundance of isobutyrate kinase (Figure IV-C in the Data Supplement) and 2-methylbutanoyl-CoA (Figure 6F) dehydrogenase in the metagenomic samples (Figure 6E-F). For 2-methylbutanoyl-CoA dehydrogenase 86% of the gene-variants were also mapped to those 60 MAGs displayed in Figure 6F (ANOVA Tukey’s HSD *p* < 0.05 for cluster 2 LPD subjects), but none of these had significant matches to isobutyrate kinase. Isobutyrate kinase was found in 86 MAGs (Figure IV-A in the Data Supplement) belonging to *Bacteroides*, Ruminococcaceae, Alistipes, Desulfovibrionaceae and Lachnospiraceae, and whose cumulative relative abundance varied (Figure IV-B in the Data Supplement) substantially between LPD clusters.

**Figure 6.**
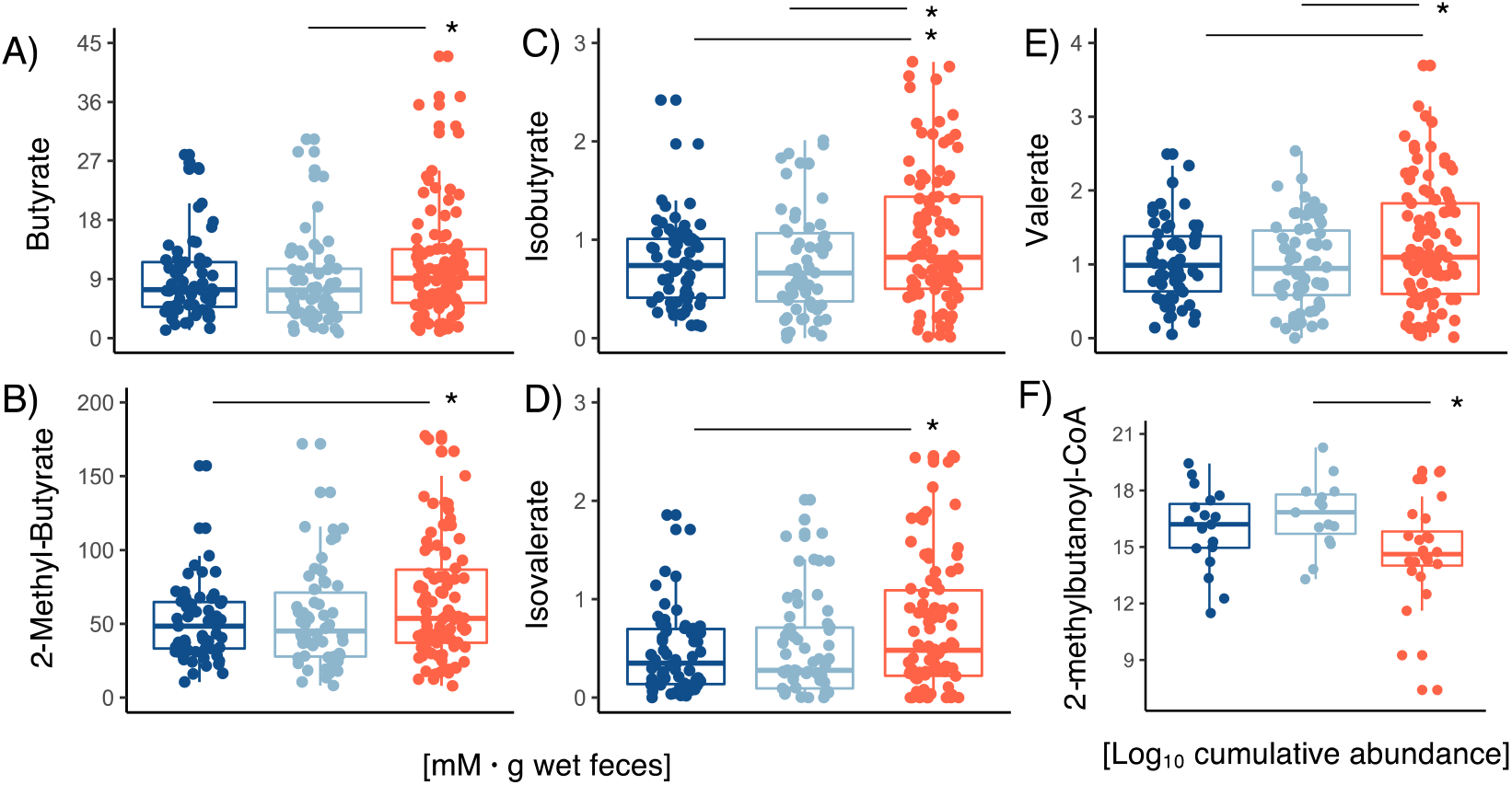
Short chain fatty acid concentrations. Range of fecal **A**) butyrate, **B**) 2-methylbutyrate, **C**) isobutyrate, **D**) isovalerate, **E**) valerate concentrations within the different LPD clusters. Cumulative abundance 2-methylbutanoyl-CoA genes screened on metagenomes within LPD clusters. Stars show statistical level of significance (\**p*≤ 0.05)

## Discussion

It is well established that certain LPD profiles are associated with elevated CVD risk, but relatively little is known on the links between GM and LPD. Building on recently published LPD profiles of 262 adult individuals ^6^ the present study investigates the correlations between LPD-profiles and GM, and its genetic functional assignments.

Stratification of study participants based on their LPD profiles yielded three LPD clusters (1A, 1B and 2) that corresponded well with within- and outside-suggested levels of total cholesterol, triglycerides, LDL, HDL and VLDL as those recommended by the NIH^40^ and as shown in Figure 2A. Our study demonstrates that lower levels of total HDL are associated with a decrease in the concentration of large subfractions (e.g. HDL2a and HDL2b), while higher levels of LDL correspond with an increase in the concentration of large LDL subfractions (e.g. LDL1). Similarly, high levels of cholesterol corresponded with high levels of circulating levels of VLDL. As confirmed by our results and others, the LPD profiles are influenced by host factors like age, sex and BMI ^5,10^ These components are able to explain up to 25% of the total variance in the LPD. To the best of our knowledge, this study represents the first to show the contribution of LPD subfractions to the collective levels of cholesterol, cholesterol-types and triglycerides, as well as recommendations among an age-/BMI-diverse group of apparently healthy adults.

Increasing evidence supports the role of GM to modulate lipids homeostasis and development of dyslipidemia ^16,42–44^. GM profiling did not show major differences in the number of sequence-variants and gene-richness counts among subjects with remarkably distinct LPD profiles (e.g., C1A, C1B and C2 clusters). Despite the fact that a bimodal distribution of gene-richness counts was reproduced as in previous studies ^45,46^ no significant differences in the gene-frequencies between normal and overweight participants were observed.

Beta diversity analyses showed significant differences that discriminated LPD clusters (e.g., Figure 4A). Lachnospiraceae members correlated positively with small LDL particles (e.g., LDL3, LDL4 and LDL5), ILDL and VLDL, while Ruminococcaceae, a subgroup of Lachnospiraceae phylotypes and other less abundant families showed positive correlations with large particles of HDL (HDL2a and HDL2b (see e.g., Figure 4G). Moreover, in agreement with our findings, a recent large-scale study published by Vojinovic et al. ^5^ also reported that Lachnospiraceae and Ruminococcaceae members were related to the HDL/LDL ratios. High HDL levels have been consistently correlated to a low risk of developing CVD ^7,8^ and recent evidence support that the heterogeneity of HDL display different associations with the incidence of CVD and metabolic syndrome ^7,47,48^. Recent findings suggest that *Akkermansia muciniphila* induces expression of low-density lipoprotein receptors and ApoE in the hepatocytes, facilitating the clearance of triglyceride-rich lipoprotein remnants, chylomicron remnants, and intermediate-density lipoproteins, from circulation ^42^. In line with this, our study elucidates a possible link between dyslipidemia and the metabolic potential of MAGs for biosynthesizing important bioactive compounds such as vitamin B complex and peptidoglycans, as well as SCFA metabolism. Among these compounds, pantothenate (Vit_B5_), Vit_B6_ and folate (Vit_B9_) have been inversely associated with low-grade inflammation ^49^ and mortality risk of CVD in a mechanism that may involve regulation of blood homocysteine concentrations ^50^ and one-carbon metabolism ^51^. SCFA like butyrate and valerate have been shown to decrease total cholesterol and the expression of mRNA associated with fatty acid synthase and sterol regulatory element binding protein 1c, to enhance mRNA expression of carnitine palmitoyltransferase-1α (CPT-1α) in liver ^52,53^, as well as to ameliorate arteriosclerosis via ABCA1-mediates cholesterol efflux in macrophages ^54^. Biosynthesis of peptidoglycans by some GM members has been associated with incidence of stenotic atherosclerotic plaques and insulin resistance ^55,56^. However, emerging evidence suggests that these potent signaling molecules play positive roles for enhancing systemic innate immunity ^57^ and neurodevelopmental processes ^58^, relaying on a species-dependent fashion ^59^. In conclusion, our study provides evidence that GM members (e.g., MAGs) and their genes related to the biosynthesis of bioactive molecules needed to carry out lipid metabolism, e.g., vitamin B complex and S/B-CFA, are more abundant among subjects with healthier LPD profiles (e.g., higher HDL2a, HDL2b, and lower LDL). Furthermore, variations in LPD subfractions correlates with differences in the GM composition ^5^, but these are not necessarily associated to a higher or lower microbial diversity as reported in previous studies ^45,46^. Given the cross-sectional nature of our study and its inherent limitations, it is not possible to depict the mechanism by which GM may influence variability in LPD subfractions. However, our results provide evidence for GM considerations in future research aiming at unravelling the processes of LPD particles assembly through longitudinal mechanistic approaches that include the activity of enzymes and transfer proteins, membrane modulators ^60^ and integrative multi-omics.

## ARTICLE INFORMATION

### Supplemental Materials

Data Supplement Figures I – IV

Supplementary-table_1

