## Supplementary Figures for "Gut microbiome and its cofactors are linked to lipoprotein distribution profiles"

### DATA SUPPLEMENT

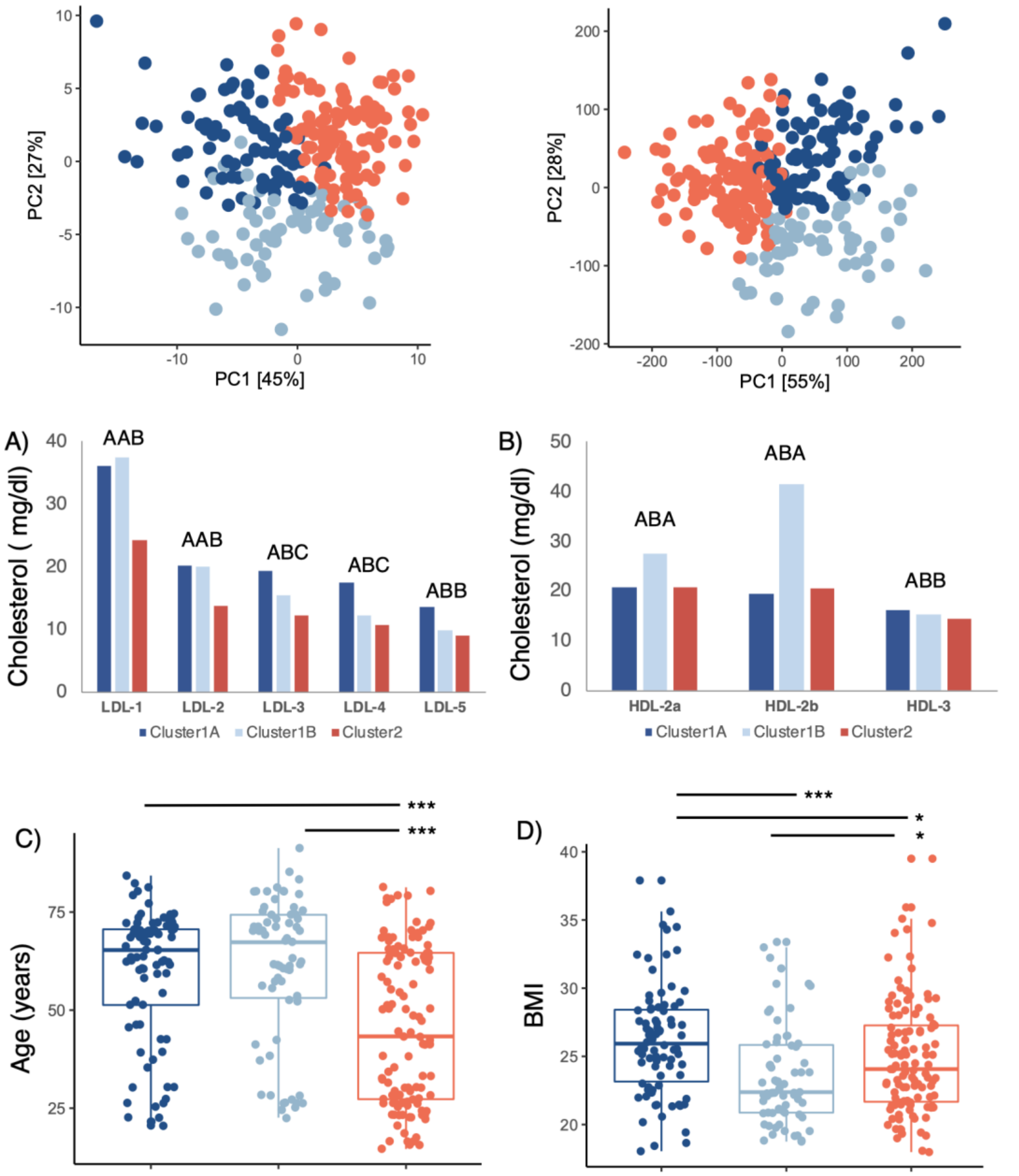


**Online Figure I. Cholesterol sub-fractions distribution and covariates adjusted by sex effect.**
Principal Component Analysis (PCA) discriminates clusters based on **A**) scaled and **B**) non-scale lipoproteins data. Average concentrations of circulating **C**) LDL and **D**) HDL sub-fractions among LPD clusters. Multiple comparisons ($P \leq 0.05$) were carried out by Tukey's HSD and display by letters A, B and C. Distribution of **E**) age and **F**) body mass index (BMI) between subjects belonging to C1A, C1B and C2 a. Stars show statistical level of significance (*$p \leq 0.05$, ***$p \leq 0.001$)

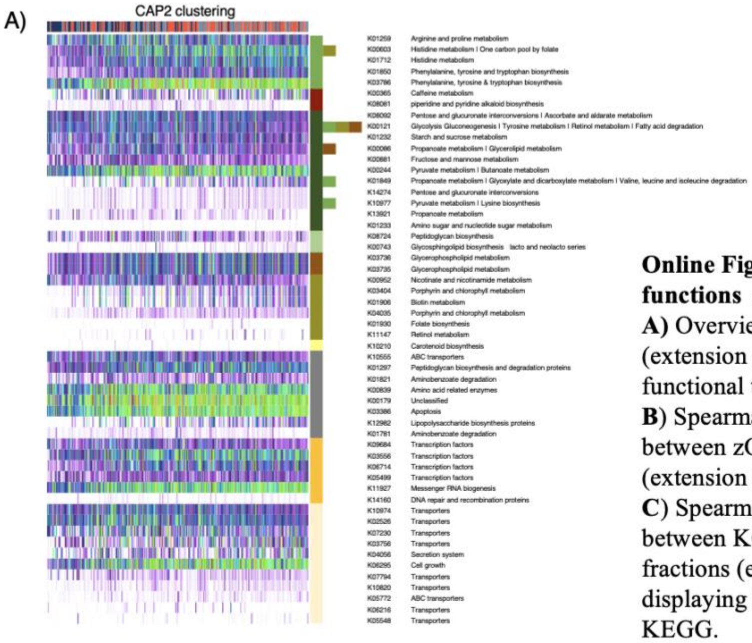


**Online Figure II. zOTUs and KO functions at 3rd level KEGG**
**A)** Overview of selected KOs-PICRUSt (extension of Fig. 2B) displaying functional traits at 3rd level KEGG.
**B)** Spearman's rank correlations between zOTUs and LPDs sub-fractions (extension of Fig. 2G).
**C)** Spearman's rank correlations between KOs-PICRUSt and LPDs sub-fractions (extension of Fig. 2H) displaying functional traits at 3rd level KEGG.

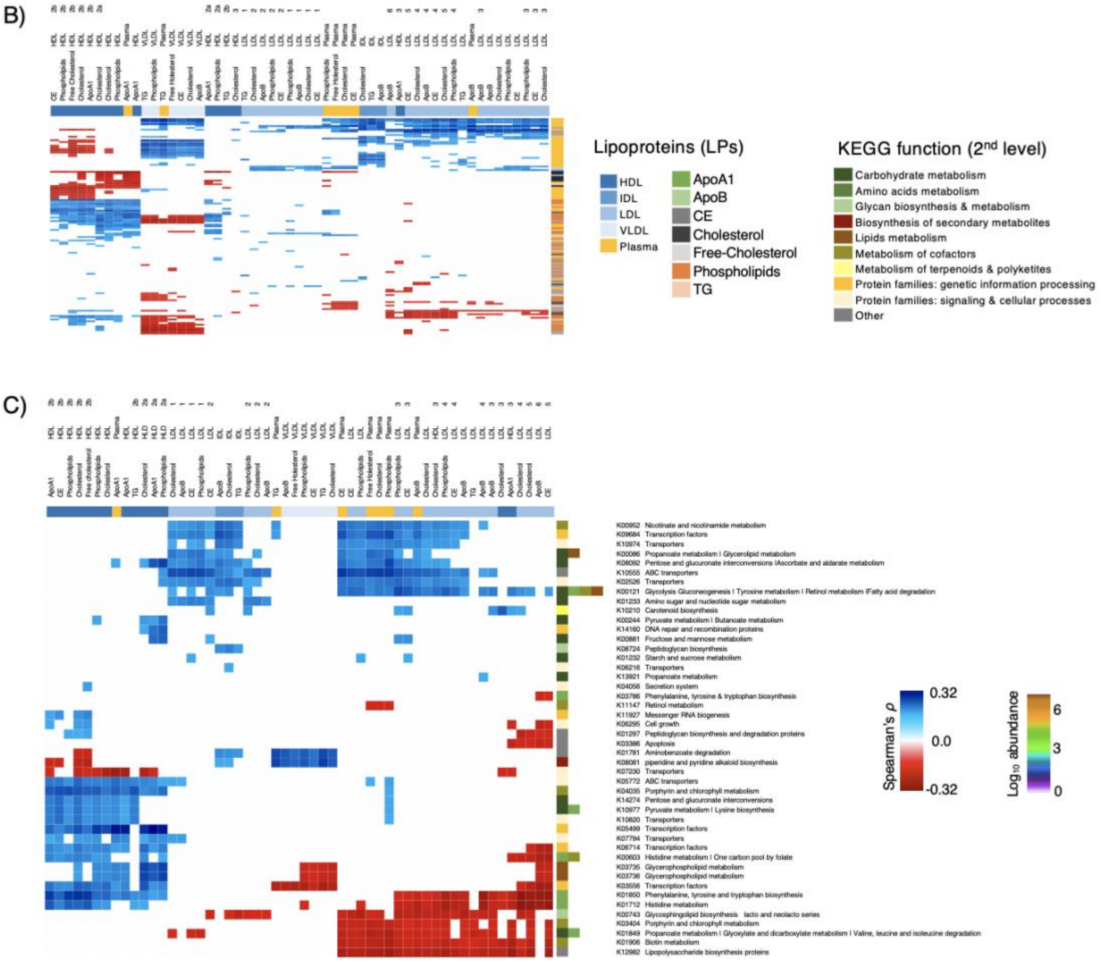

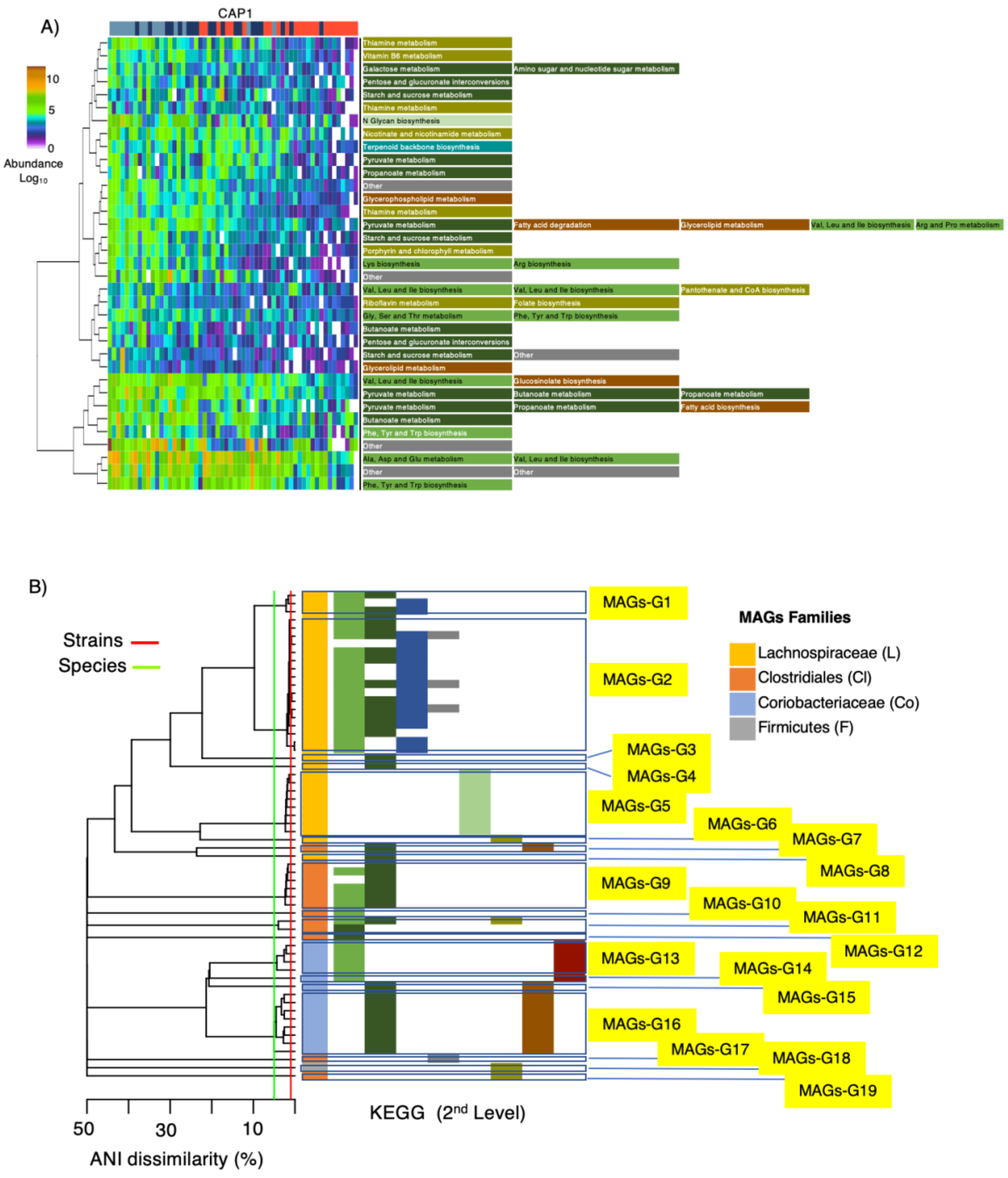


**Online Figure III**
**A**) Overview of most discriminatory (extension of Fig. 5B) KOs summarized to their 3rd level KEGG function.
**B**) Phylogeny of MAGs outlined into species groups (95 ANI) – (extension of Fig. 5D)

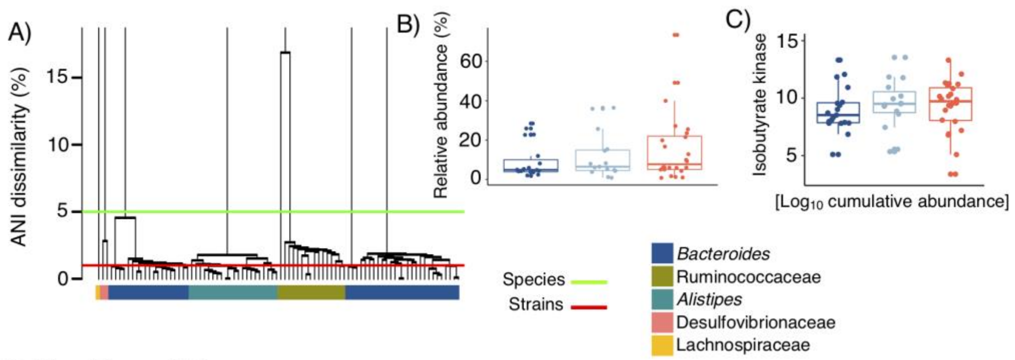


**Online Figure IV**

A) Phylogeny of MAGs outlined into species groups (95 ANI) containing isobutyrate kinase genes.
B) Cumulative relative distribution MAGs within LPDs clusters.
C) Abundance of branched-chain fatty acid (BCFA) genes kinase within LPDs clusters.
